# Pathology of naturally acquired high pathogenicity avian influenza virus H5N1 infection in seabirds

**DOI:** 10.1101/2023.02.17.528990

**Authors:** Fabian ZX Lean, Marco Falchieri, Natalia Furman, Glen Tyler, Caroline Robinson, Paul Holmes, Scott M Reid, Ashley C Banyard, Ian H Brown, Catherine Man, Alejandro Núñez

**Author notes:** Corresponding author: Fabian Lean.

## Abstract

The re-emergence of the high pathogenicity avian influenza virus (HPAIV) subtype H5N1 in the United Kingdom in 2021-2022 has caused unprecedented epizootic events in wild birds and poultry. During the summer of 2022 there was a shift in virus transmission dynamics resulting in increased HPAIV infection in seabirds and consequently a profound impact on seabird populations. To understand the pathological impact of HPAIV in seabirds, we have evaluated the virus distribution and associated pathological changes in the tissues of great skua (*Stercorarius skua*, n=8), long tailed skua (*Stercorarius longicaudus*, n=1), European herring gull (*Larus argentatus*, n=5), and black-headed gull (*Chroicocephalus ridibundus*, n=4). Grossly there was gizzard ulceration in one great skua and pancreatic necrosis in four herring gulls, which were confirmed for virus infection *in situ* by immunohistochemistry. Microscopical analysis revealed neuro-, pneumo-, lymphoidand cardiotropism of HPAIV H5N1, with the most common virus-associated pathological changes being pancreatic and splenic necrosis. Examination of the reproductive tract of the great skua revealed HPAIV-associated oophoritis and salpingitis, and virus replication within the oviductal epithelium. Across the birds, epitheliotropism was evident in the intestine, nasal turbinate, and trachea. This was, in contrast, not observed in the 2021 summer mortality event in great skuas and may be significant for the disease epidemiology observed in 2022. The emergence of HPAIV in seabirds, particularly during summer 2022, has challenged the dogma of HPAIV dynamics, posing a significant threat to wild bird life with potential implications to the reproductive performance of seabirds of conservation importance.

## Introduction

High pathogenicity avian influenza virus (HPAIV) H5N1 clade 2.3.4.4b has reemerged in Europe and the United Kingdom (UK) during 2020-2021 and 2021-2022 seasons (defined as start of each October) and has brought about a series of epizootic events in poultry and wild birds. The re-emergence of HPAIV H5N1 clade 2.3.4.4.b in Europe and the UK during 2021-2022 has also contributed to the transAtlantic dissemination of virus into North America likely mediated though migratory wild birds.^12^

Conventionally it is understood that Anseriformes are the carrier for HPAIV during the winter period in Western Europe. However during the 2021-2022 HPAIV season in the UK, there was a shift in infection from Anseriformes predominating in the colder months to a series of explosive outbreaks in seabird species across the northern coast of Scotland during summer.^4^ During summer 2021, infection with H5N1 was detected in great skuas^6^ but those events, alongside sporadic small-scale outbreaks across northern Europe were the only cases of H5N1 reported during the summer months. In contrast during summer 2022, infection in great skuas (*Stercorarius skua*) was detected several months earlier than seen during 2021 and was followed by extensive outbreaks in a number of shorebird species (Order Charadriiformes).^16^ High mortality events in seabirds including northern gannet, great skua and several species of gull species were observed.^4,16^ Seabirds from the Laridae family have been previously associated with infection with low pathogenicity avian influenza virus (LPAIV). ^18,20,24,32,50^ However, a recent experimental model has demonstrated previous exposure of the European herring gulls with LPAIV H5N1 or H13N6 only confer partial protection to subsequent HPAIV H5N8 clade 2.3.4.4b challenge.^46^

Prior to the unusual increased in cases during summer 2022, HPAIV-associated disease in the Laridae has been sporadically reported in East Asia and Europe, often in small numbers.^1,2,12,15,34,36,37,39^ More recently there has been an increased detection of HPAI positive seabirds or Charadriiformes,^16,27^ and critically, mortality events in seabirds associated with HPAIV infection reported in the UK, Europe and North America.^5,6,44^ Data collected through the avian influenza wild bird passive surveillance in Great Britain have shown a rise in high pathogenicity H5Nx positive birds within the *Laridae* family from 1.3% during the 2020-2021 season and up to 15% within the 2021-2022 season. The potential maintenance of HPAIV in seabirds introduces further uncertainty on the transmission dynamics at both the local and global levels.

One of the hypotheses for the enzootic transmission of HPAIV in wild birds in Europe is the maintenance in wild birds during summer in Northern Europe.^41^ Previously Anseriformes were thought to be responsible for transmission given potential virus adaptation in the host^10,11^ but the expansion of avian taxa susceptibility for HPAIV and increased incidence of disease also challenges the status quo.

Here we evaluate the relationship between virus antigen distribution in tissues and the associated pathological changes in great skua (*Stercorarius skua*), long tailed skua (*Stercorarius longicaudus*), European herring gull (*Larus argentatus*), and black-headed gull (*Chroicocephalus ridibundus*) which succumbed to natural infection of HPAIV during summer of 2022 and comment upon the potential for these birds as reservoirs of infection.

## Materials and methods

### Post-mortem examination

Carcasses received at Scotland’s Rural College, NatureScot or APHA regional laboratories were frozen for transport, and thawed for necropsy at APHA Weybridge. The herring gulls came from wildlife rehabilitation centres (Sussex and Cornwall), whereas other samples were retrieved from birds that have been submitted having being found dead in the wild. The great skuas carcasses originated from colonies on Shetland including Scatness (Mainland), Noss (Island), Noness (Mainland), and long tailed skua carcass originated from Clumlie (Mainland). Oropharnygeal (OP) and cloacal (C) swabs were tested to confirm infection status with HPAIV H5N1 by standard tests as described previously.^35^ Major organs were fixed in 10% neutral buffered formalin for microscopic evaluation.

### Histopathology and immunohistochemistry

Formalin fixed tissues samples were processed using routine histological methods into paraffin blocks. Tissues were sectioned at 4 μm thickness and stained with hematoxylin and eosin (H&E) for histological evaluation and immunohistochemical labelling using a monoclonal antibody against the nucleoprotein (NP) of influenza A virus for the detection of influenza viral antigen as described previously.^38^ The tissues were assessed on conventional light microscope for histopathology: Absent -, minimal +, mild ++, moderate +++, severe ++++; and abundance of virus antigens: Absent -, rare +, scattered ++, confluent +++, abundant ++++.^6^

## Results

### Post-mortem findings

The clinical disease reported in the captive herring gulls included cyanotic heads, gasping, muscle twitching, diarrhoea, and sudden deaths. Other birds obtained from the wild were found dead.

All submitted birds were in good body condition. On necropsy, the great skuas (n=8, 6 females and 2 males) were moderately autolysed. Only one of the birds had multifocal, approximately 1 to 2mm diameter, red to brown ulcers at the proventricular-gizzard junction (**Figure 1a**). The long tailed skua (n=1, male) was moderately autolysed and otherwise unremarkable. For the herring gulls (n=5, 2 male and 3 gender not determined) were mildly autolysed. Post-mortem examination findings included multifocal faint tan areas in the pancreatic parenchyma (n=4; **Figure 1b**), suggestive of necrosis, mild splenomegaly (n=3), and intestinal nematodiasis (n=1), A black headed gull (n=1, 2 male and 2 gender not determined) was examined but was unremarkable and with severe autolysis.

**Figure 1.**
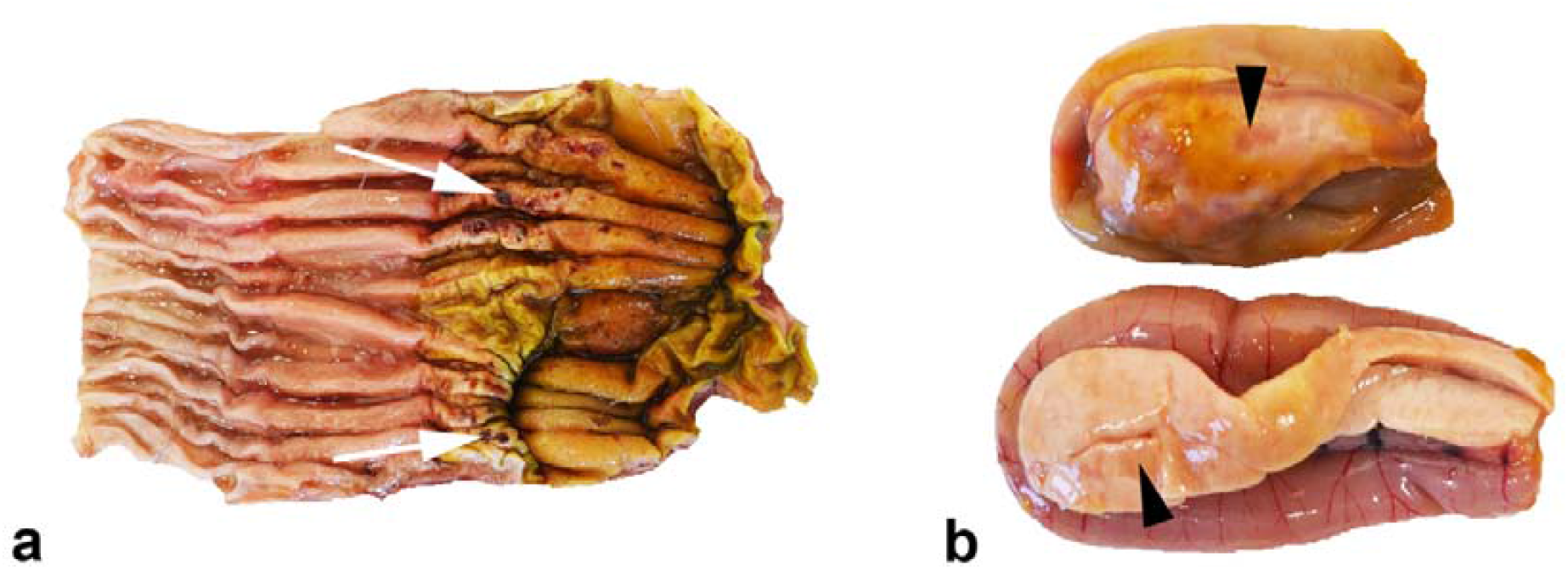
Gross lesions of HPAIV H5N1 infected seabirds. Multifocal areas of dark red depression (white arrow) on the mucosa of the proventricular-gizzard junction (a), great skua (*Stercorarius skua*). Multifocal to coalescing pale tan areas in the pancreas consistent with necrosis (b), European herring gull (*Larus argentatus*).

### Histopathology and viral immunohistochemistry

In the great skuas, virus antigen was consistently detected in the heart, brain, kidney, lung, and pancreas of all birds examined (**Table 1**). The pancreas was particularly severely affected, with moderate to severe areas of confluent necrosis. These necrotic areas correlated with moderate to abundant distribution of virus antigens in all skuas examined (**Figure 2a**; n=8/8). Correlative viral IHC and histology also revealed viral-associated myocardial necrosis (**Figure 2b**, n=2/8), splenic necrosis (**Figure 2c**; n=3/4), and renal tubular necrosis (**Figure 2d**; n=5/8). In the proventriculus of a great skua where ulceration was noted at post-mortem (**Figure 1a**), histological and IHC examination confirmed immunolabelling of the mucosa (**Figure 3a**) and glandular epithelium (**Figure 3b**). The mucosal damage was extensive and was replaced with necrotic cellular debris, degenerated heterophils and fibrin deposition (**Figure 3a, b**). Similar proventricular mucosa damage was also observed histologically in other two great skuas where lesions were not observed grossly. Nevertheless, viral immunolabelling in the proventriculus (n=6/8) and gizzard (n=5/8) were more common than histopathological changes (n=1, 3/8; respectively). Only one long tailed skua was examined, which revealed severe pancreatic necrosis and mild splenic necrosis (**Table 1**).

**Table 1.** Summary of histopathology and viral immunohistochemistry findings.

| Tissue | Great Skua (n=8)<br>*Stercorarius skua* |  |  |  | Long tailed skua (n=1)<br>*Stercorarius longicaudus* |  |  |  | European herring gull (n=5)<br>*Larus argentatus* |  |  |  | Black-headed gull (n=4)<br>*Chroicocephalus ridibundus* |  |  |  |
| --- | --- | --- | --- | --- | --- | --- | --- | --- | --- | --- | --- | --- | --- | --- | --- | --- |
|  | Histopathology<sup>a</sup> |  | IHC<sup>b</sup> |  | Histopathology |  | IHC |  | Histopathology |  | IHC |  | Histopathology |  | IHC |  |
|  | n (%) | Grade | n (%) | Grade | n (%) | Grade | n (%) | Grade | n (%) | Grade | n (%) | Grade | n (%) | Grade | n (%) | Grade |
| Skin | 0/8 (0) | - | 5/8 (63) | + | 0/1 (0) | - | 1/1 (100) | + | 0/4 (0) | - | 0/4 (0) | - | 0/4 (0) | - | 1/4 (25) | + |
| Skeletal muscle | 0/8 (0) | - | 7/8 (88) | + | 0/1 (0) | - | 1/1 (100) | ++ | 0/5 (0) | - | 5/5 (100) | + | 1/3 (33) | + | 3/3 (100) | + to ++ |
| Heart | 2/8 (25) | + | 8/8 (100) | + to +++ | 0/1 (0) | - | 1/1 (100) | ++++ | 1/5 (20) | ++ | 5/5 (100) | + to ++ | 0/4 (0) | - | 4/4 (100) | + to ++++ |
| Brain | 3/8 (38) | + to ++ | 8/8 (100) | + to ++++ | 0/1 (0) | - | 1/1 (100) | +++ | 4/5 (80) | ++ | 5/5 (100) | ++ to ++++ | 1/4 (25) | + | 4/4 (100) | ++ to ++++ |
| Spleen | 3/4 (75) | ++ | 3/4 (75) | ++ to ++++ | 1/1 (100) | ++ | 1/1 (100) | ++++ | 4/5 (80) | + | 5/5 (100) | + to ++ | n/a | n/a | n/a | n/a |
| Kidney | 5/8 (63) | + | 8/8 (100) | + to ++ | 0/1 (0) | - | 1/1 (100) | ++++ | 0/5 (0) | - | 3/5 (60) | + | 1/1 (100) | + | 1/1 (100) | ++ |
| Nasal turbinate | n/a | n/a | n/a | n/a | n/a | n/a | n/a | n/a | 4/4 (100) | + to ++++ | 3/4 (75) | ++ to +++ | n/a | n/a | n/a | n/a |
| Trachea | 0/8 (0) | - | 5/8 (63) | 1 to ++ | 0/1 (0) | - | 1/1 (100) | ++ | 3/3 (100) | + to ++ | 3/3 (100) | + | n/a | n/a | n/a | n/a |
| Lung | 1/8 (13) | + | 8/8 (100) | + to ++++ | 1/1 (100) | + | 1/1 (100) | ++++ | 3/5 (60) | + | 5/5 (60) | + to ++++ | 1/2 (50) | +++ | 2/2 (100) | ++++ |
| Proventriculus | 3/8 (38) | + to +++ | 6/8 (75) | + to ++ | n/a | n/a | n/a | n/a | 0/5 (0) | - | 1/5 (20) | + | 0/4 (0) | - | 2/4 (50) | + to ++ |
| Gizzard | 1/8 (13) | ++ | 5/8 (63) | + to ++ | 0 | ++ | 1/1 (100) | ++ | 1/5 (20) | + | 1/5 (20) | ++ | 0/3 (0) | - | 3/3 (100) | + |
| Liver | 2/8 (25) | + to ++ | 7/8 (88) | + to ++ | 1/1 (100) | + | 1/1 (100) | ++++ | 3/5 (60) | + to ++ | 4/5 (80) | + to ++ | 1/3 (33) | + | 1/3 (33) | ++ |
| Pancreas | 8/8 (100) | +++ to ++++ | 8/8 (100) | +++ to ++++ | 1/1 (100) | ++++ | 1/1 (100) | +++ | 5/5 (100) | + to ++++ | 5/5 (100) | ++ to ++++ | n/a | n/a | n/a | n/a |
| Duodenum | 1/8 (13) | + | 5/8 (63) | + to ++ | n/a | n/a | n/a | n/a | 2/5 (40) | + | 3/5 (60) | + to ++ | n/a | n/a | n/a | n/a |
| Ovary | 6/6 (100) | + to +++ | 6/6 (100) | + to ++++ | n/a | n/a | n/a | n/a | n/a | n/a | n/a | n/a | 0/4 (0) | - | 4/4 (100) | + |
| Oviduct | 5/5 (100) | + to +++ | 5/5 (100) | ++ to +++ | n/a | n/a | n/a | n/a | n/a | n/a | n/a | n/a | n/a | n/a | n/a | n/a |
| Testis | 0/2 (0) | - | 0/2 (0) | - | 0/1 (0) | - | 1/1 (100) | ++ | n/a | n/a | n/a | n/a | n/a | n/a | n/a | n/a |

<sup>a</sup> Histopathology grade: Absent -, minimal +, mild ++, moderate +++, severe ++++
<sup>b</sup> Immunohistochemical grade: Absent -, rare +, scattered ++, confluent +++, abundant ++++

**Figure 2.**
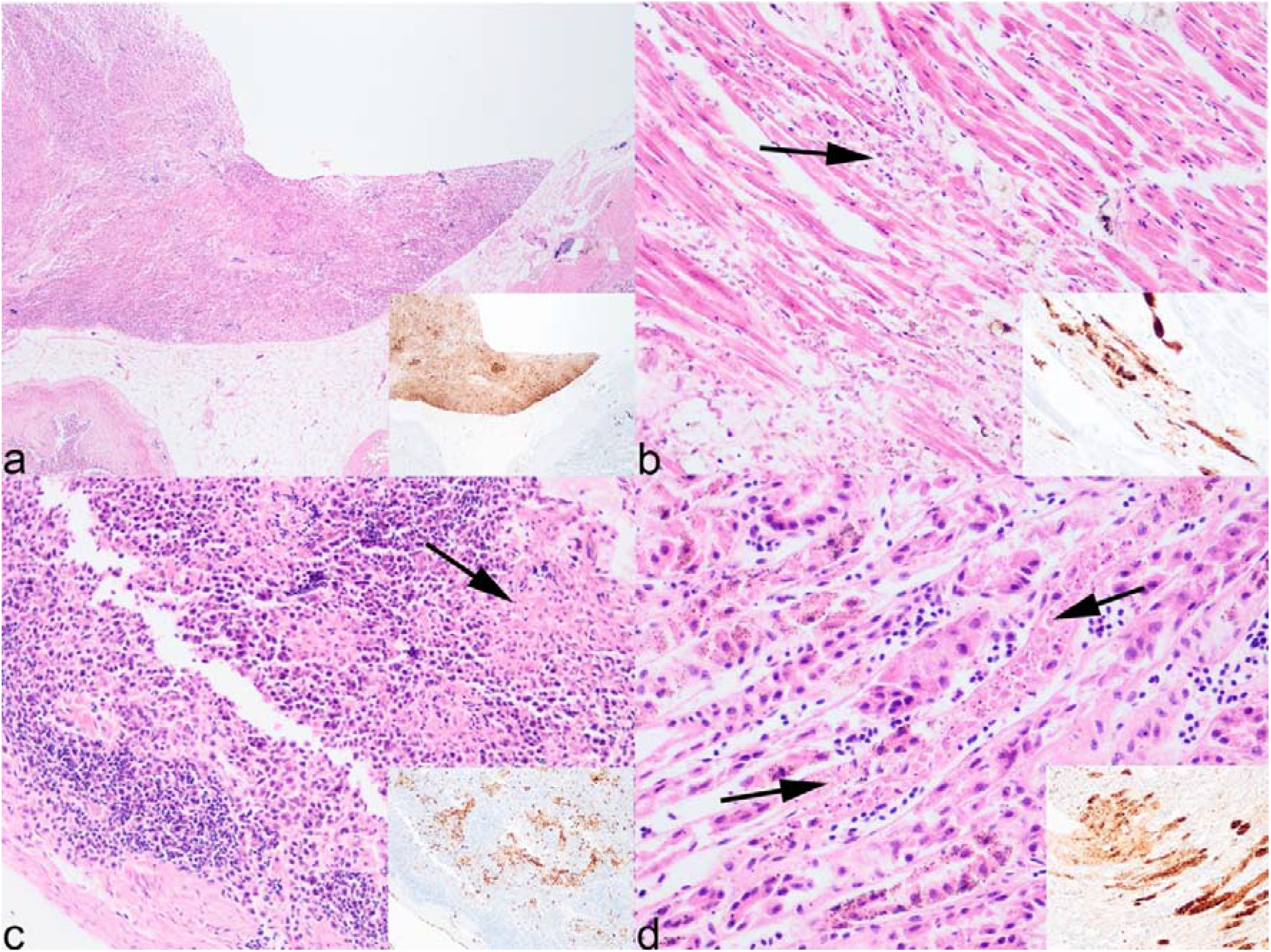
Microscopic findings of great skua (*Stercorarius skua*) infected with HPAIV H5N1. Severe, confluent, pancreatic necrosis (a). Moderate, multifocal, myocardial necrosis (b). Mild, multifocal, splenic necrosis (c). Mild, multifocal, renal tubular necrosis (d). Co-localisation of virus antigens with areas of necrosis in various tissues (a-d, insets). Arrows indicate area of necrosis. Histological images were taken at 40x (a), 200x (d) and 400x (b, c) and immunohistochemical insets were taken at 40x (a), 200x (b, c) and 400x (d).

**Figure 3.**
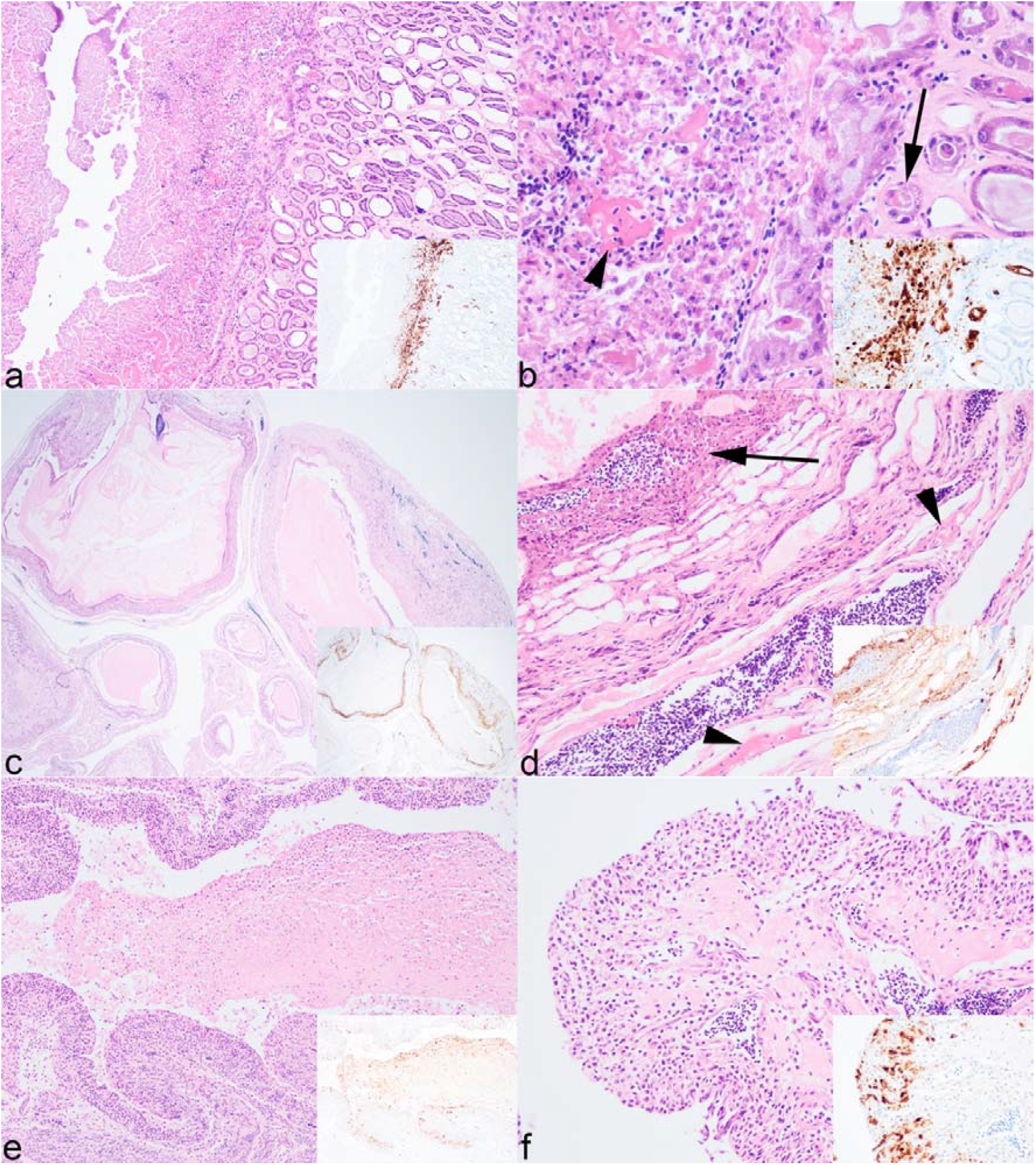
Microscopic findings of great skua (*Stercorarius skua*) infected with HPAIV H5N1. (a and b) Moderate, focal, gizzard necrosis and fracturing of koilin (a), with evidence of mucosa epithelial degeneration (arrow), and deposition of cellular debris, degenerated heterophils, extravasated erythrocytes, and fibrin (arrowhead) within disrupted koilin layer (b). Moderate, diffuse, oophoritis (c) characterised by necrosis of theca interna (arrow) and stroma, and with fibrin deposition within the mural typically around blood vessels (d, arrowhead). Moderate necrotising salpingitis, with abundant intra-luminal debris (e) and mucosa is eroded and infiltrated with heterophils and lymphocytes (f). Co-localisation of virus antigens with areas of necrosis in various tissues (a-f) and intra-luminal debris in the oviduct (e) (insets). Arrows indicate area of necrosis. Histopathology images were taken at 20x (c), 100x (e), 200x (a, d, f), and 400x (b) magnifications and immunohistochemical insets were taken at 20x (c), 100x (a, e), 200x (d) and 400x (b, f) magnifications.

The reproductive tract was only available for examination from the great skuas. Virus antigens were detected in the ovaries (n=6/6) and oviducts (n=5/5) but not in the testis (n=0/2). In the ovaries, there was confluent distribution of viral antigens (**Figure 3c**), being predominantly present in the theca interna and occasionally transmural of the pre-ovulatory follicles. This was associated with necrosis within the tunica interna and blood vessels of the stroma, and the stromal wall was moderately to markedly expanded with lymphoplasmacytic cells and fibrin deposits (**Figure 3d**). In the oviduct, there was a moderate amount of intra-luminal debris, ulcerated mucosa, and heterophilic and lymphocytic infiltration of submucosa wall observed. Virus antigens were present in the debris (**Figure 3e**), remaining mucosa epithelium and submucosal cells (**Figure 3f**).

In the herring gulls, viral antigen was found in the brain (n=4/5), lung (n=5/5), pancreas (n=5/5), nasal turbinate (n=3/4). Lesions that were consistently associated with viral antigens include the pancreas (n=5/5), brain (n=4/5) and nasal turbinate (n=3/4). Pancreas necrosis was multifocal to confluent acinar necrosis (**Figure 4a**). In the spleen mild lymphoid depletion was observed (**Figure 4b**). In the brains, there was mild neuronal necrosis and dispersed degenerated heterophils within the neuropil, and in the cerebellum, there was occasional loss of Purkinje cells attributed to viral infection (**Figure 4c**). In the lungs, there were mild to moderate air capillary necrosis and occasionally fibrin deposition in air capillary walls. Rhinitis ranged from mild changes including scant neutrophilic exudate with occasional intra-epithelial neutrophils, or in severe damage with abundant exudation, complete loss of mucosa with submucosa necrosis and fibrins deposition (**Figure 4d**). Incidental findings included presence of intestinal cestode and proventricular nematode in two separate herring gulls but not associated with overt intestinal pathology.

**Figure 4.**
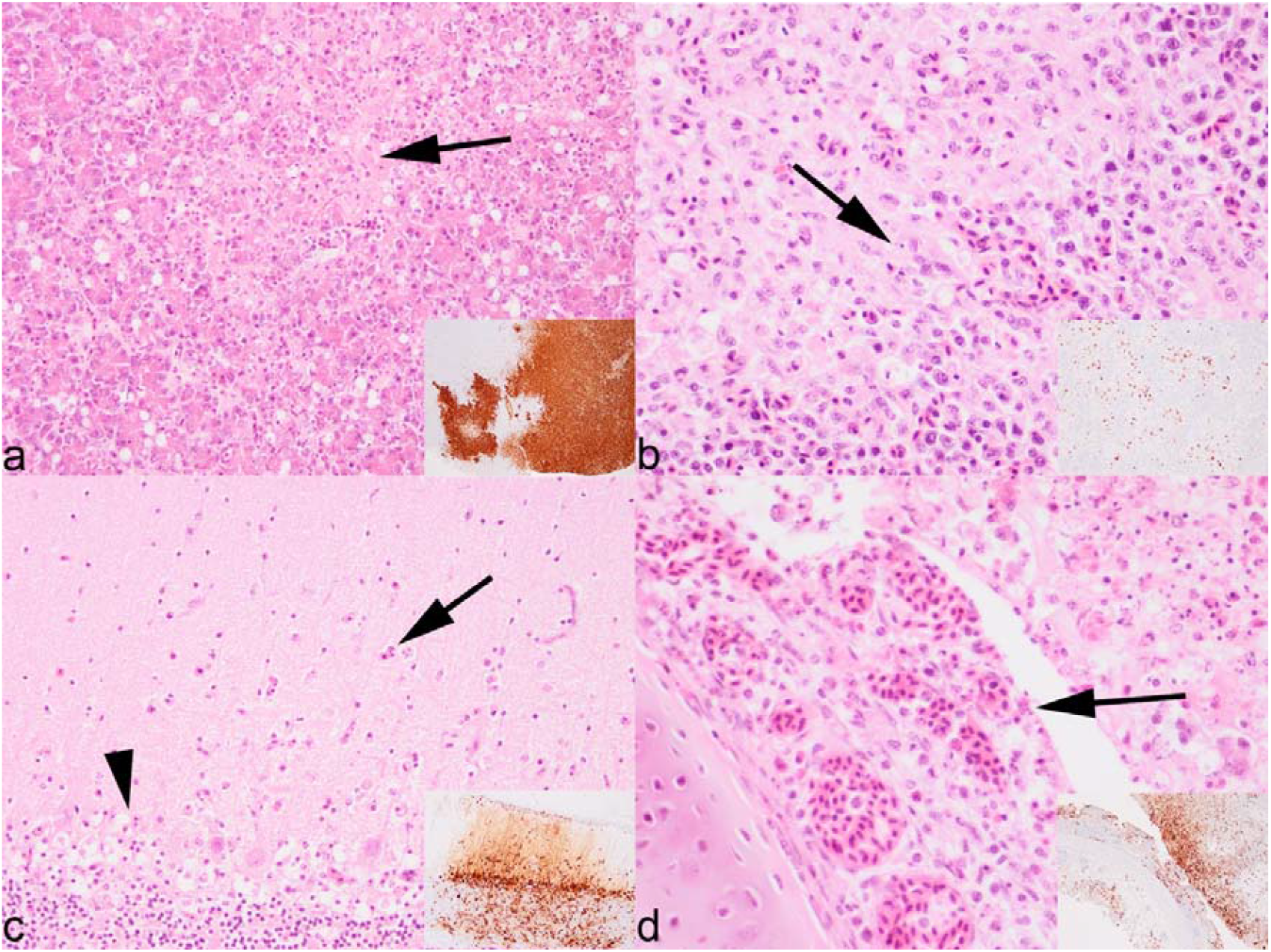
Microscopic findings of European herring gull (*Larus argentatus*) infected with HPAIV H5N1. Mild, multifocal, necrosis and vacuolar degeneration of the pancreatic acinar cells (a). Mild, multifocal necrosis of splenic white pulp (b). Minimal, multifocal, neuronal necrosis with scattered heterophils in neuropil (arrow) and loss of Purkinje cells, cerebellum (arrowhead) (c). Severe, confluent, necrotising rhinitis with extensive loss of epithelial layer (arrow), abundant exudation and infiltration of heterophils and lymphocytes in submucosa (d). Histopathology images were taken at 200x (a, c) and 400x HE (b, d) magnifications and immunohistochemical insets at 40x IHC (a) and 100x IHC (b, c, d) magnifications.

## Discussion

During the 2020-2021 and 2021-2022 outbreaks of high pathogenicity avian influenza in the UK, there has been an increased detection of HPAIV H5N1 in seabirds of the Orders Suliformes and Laridae.^3,6,16,34^ This investigation of naturally acquired infection revealed that gross pathology was limited to pancreatic necrosis in the herring gull and proventricular ulceration in the great skua. The pancreatic changes were less characteristic compared to that in Galliformes and Anseriformes and required immunohistochemical confirmation. Microscopic evaluation confirmed a multi-systemic HPAIV infection including neuro-, cardio, and pneumo-tropism, which may have been contributary to the mortalities seen. In addition, acute reproductive damage in female great skuas’ was noted. Overall, seabirds are highly susceptible to developing pathologies in multiple organ systems following HPAIV infection.

Historically, gull species from the Order Laridae have been associated with infection with LPAIV including H11, H13 and H16 subtypes.^24,32,50^ These infections have been predominantly associated with replication in epithelial cells of intestine that has been hypothesised to facilitate faecal-oral transmission in black headed gulls.^25^ For HPAIV, natural infection has been reported in great skuas, European herring gulls, black-headed gulls, and great black-backed gulls (*Larus marinus*).^1,2,6,12,15,34,37,39^ The most common and severe pathology in all birds examined was pancreatic necrosis associated with viral infection (except for in the black-headed gull where the pancreas was unavailable), followed by splenic necrosis and pneumonia (except in the long tailed skua). Such lesions are like those reported from experimentally challenged common gulls with H5N1, plus naturally infected black headed gulls and herring gulls, together with a recent report of naturally infected sandwich terns (*Thalasseus sandvicensis*).^9,23,43,44,46^ Although RT-PCR was not conducted on tissues from the current investigation, the abundance of virus antigens in the heart, brain, kidney, spleen, lung, pancreas, and liver strongly suggests the utility of these organs for diagnostic evaluation.

It has been proposed that the enterotropic adaptation of HPAIV in wild waterbirds has facilitated long term persistence and dissemination in these species with virus being maintained in stable equilibrium without undue pathological impact on the host.^10,11^ In this report, we noted higher level of immunolabelling in the trachea, proventriculus, gizzard and duodenum in the great skua, and additionally antigens detected in the respiratory and enteric epithelium, which were generally absent in the previous GB epizootic in great skua in 2021.^6^ Similar respiratory and enterotropism was observed in the long tailed skua, herring gull and black-headed gull in this study. The nasal turbinates were only examined in the herring gull which showed viral associated rhinitis and epitheliotropism. Previously, infection of the intestine had only been observed rarely with H5N1 clade 2.2 viruses in common gulls^23^ and a laughing gull infected with an ‘Eurasian-lineage’ of H5N1.^8^ Avian influenza viruses preferentially bind the α-2,3 sialic acid residues.^13^ Based on lectin histochemistry on other gull species including American herring gulls (*Larus smithsonianus*), laughing gulls (*Leucophaeus atricilla*) and ring-billed gulls (*Larus delawarensis*), the respiratory epithelium express both α-2,3 and α-2,6 sialic acids, whereas the intestinal tracts express predominantly α-2,3 and rarely α-2,6 sialic acids.^19^

The pathway of incursion in free-ranging seabirds is not understood but has been proposed to be either independent incursion or onward introductions from species movements between colonies and the movement of seabirds between mainland and islands particularly during the breeding season.^16,44^ Herring gull and great skua can opportunistically predate or scavenge on other birds,^16,26,31,51^ and this was observed in the outbreak in gannet colonies. Further, contact transmission between common gulls (*Larus canus*) and European herring gulls have been documented previously with experimental infection with HPAIV H5N1 clade 2.2 and H5N8 clade 2.3.4.4b.^23,46^ More recent HPAIV H5N1 outbreaks (June and August 2022) in wild bird rescue centres / hospitals in England (East Sussex and Cornwall) have been confirmed in herring gulls. After epidemiological assessment, the most likely source of infection appeared to be the introduction within the premises of infected / diseased herring gulls which had then transmitted the disease to the resident gulls of the same species within and among enclosures (APHA, unpublished data). It is not uncommon for skuas or gulls to congregate in high densities during the breeding season for nesting, feeding and bathing facilitate close contact. These behaviours could facilitate dissemination of HPAIV especially if virus replication is prominent in respiratory and intestinal tracts.^16^ Infections through such contact can lead to birds from other colonies becoming exposed and infected, which then themselves spread virus to new localities and susceptible avian species. Further, these seabirds are often in areas with high seal populations plus other scavenging mammals that can predate on sick or dead birds, and result in exposures of other host population types to infectious materials either directly or indirectly through the environment.^17^

The distribution and ecology of seabird populations also challenge the current understanding of HPAIV transmission at a global level. Both long tailed skua and great skua are transitory migrant birds - long tailed skuas are a passage migrant in the UK and breed in Arctic region,^22^ whereas great skuas migrate to the northernmost isles of the UK in summer for breeding and return to the coasts of Spain and Africa, and as far as Brazilian and Argentinian coasts for wintering.^14,40^ Black-headed gulls are found across the UK,^28^ and herring gulls are found throughout the year around the UK coastline and inland around rubbish tips, fields, large reservoirs, and lakes, especially during the winter months.^30^ Recent ring-recovery data revealed that great skua, European herring gulls and black-headed gulls migrate between Europe to Iceland and other North Atlantic islands, and to North America.^12^ The pelagic and migratory nature of gulls have led to suggestion of intercontinental dissemination and shaping of influenza A virus evolution.^21,24,42,48,52^

Apart from the increased mortality in seabirds during 2022 which has resulted in an immediate impact upon populations, there is generally a significant deficit in knowledge on the impact of infectious diseases on population structures across these species. However, a trend towards a reduction in breeding abundance in the UK for herring gulls, black-headed gulls and great skuas has been noted.^28-30^ The pathogenic mechanism of HPAIV on reproductive organs of wild bird is poorly documented. Previous reports have demonstrated epithelial labelling of virus antigen in the oviduct of common buzzards and peregrine falcons infected with HPAIV.^49^ In domestic poultry, both HPAIV and LPAIV infection can lead to short to long term reduction in egg production or embryonic death because of viral-induced pathology on the ovaries, oviduct, or conceptus.^7,33,45,47^ There has been an increased detection of reproductive pathologies in laying poultry, both Galliformes and Anseriformes, during the 2022 epizootic season in the UK which can be attributed to virus infection *in situ* (Lean F, unpublished). However, the impact on the poultry sector, where an abundance of eggs is produced daily, cannot be compared to seasonal reproductive cycle in seabirds and as such the longer-term impact on population densities for these species will require monitoring to assess population recovery.

In conclusion we demonstrate the susceptibility and pathology of a subset of Laridae and Suliformes following a naturally acquired infection with HPAIV H5N1 clade 2.3.4.4b. We associate rapid mortality with the observed multisystemic dissemination of viral antigen and resultant tissue damage. Reproductive pathology is also noted amongst the female great skua but the longer-term impact on population fecundity warrants further investigation.

## Acknowledgements

The authors would like to thank the scientists at APHA for their laboratory work, and colleagues at Scotland’s Rural College and NatureScot for the support.

## Ethical statement

No ethical approval was required as carcass and tissue were derived from diagnostic investigations.

## Author contribution statement

F.Z.X.L., M.F., N.F., G.T., C.R., P.H., C.M. involved in conceptualisation of the investigations. F.Z.X.L., N.F. performed the necropsies. F.Z.X.L. conducted formal analysis. A.N., A.C.B., S.M.R., I.H.B., C.M. provided project leadership, financial, and laboratory resources. F.Z.X.L. wrote the original draft. All authors reviewed and edited the manuscript.

